# Integrating human iPSC-derived macrophage progenitors into retinal organoids to generate a mature retinal microglial niche

**DOI:** 10.1101/2022.12.23.521829

**Authors:** Ayumi Usui-Ouchi, Sarah Giles, Yasuo Ouchi, Elizabeth A Mills, Martin Friedlander, Kevin T Eade

## Abstract

In the retina, microglia are resident immune cells that are essential for retinal development and function. Retinal microglia play a central role in mediating pathological degeneration in diseases such as glaucoma, retinitis pigmentosa, age-related neurodegeneration, ischemic retinopathy and diabetic retinopathy. Current models of mature human retinal organoids (ROs) derived from iPS cell (hiPSC) do not contain resident microglia integrated into retinal layers. Increasing cellular diversity in ROs by including resident microglia would more accurately represent the native retina and better model diseases in which microglia play a key role. In this study, we develop a new 3D *in vitro* tissue model of microglia-containing retinal organoids by co-culturing ROs and hiPSC-derived macrophage precursor cells (MPCs). We optimized the parameters for successful integration of MPCs into retinal organoids. We then reproducibly integrate MPCs into ROs where they develop into mature microglia (iMG) as seen by 1) migration to the appropriate anatomical locations; 2) development of a mature resting morphology; and 3) expression of mature microglial markers. We show that while in the ROs, MPCs migrate to the equivalent of the outer plexiform layer where retinal microglia cells reside in healthy retinal tissue. While there, they develop a mature morphology characterized by small cell bodies and long branching processes which is only observed *in vivo*. During this maturation process these microglia cycle through an activated phase followed by a stable mature phase characterized by cell-type specific microglia markers Tmem119 and P2ry12. This co-culture system may be useful for understanding the pathogenesis of retinal diseases involving retinal microglia and for drug discovery.

## Introduction

Microglia are resident immune cells derived from primitive hematopoietic progenitors originating from the yolk sac early in development (1). They migrate into the central nervous system, including the retina, where they mature and self-renew locally (2). Microglia play significant functional roles in the retina during development and adulthood in both normal and pathological conditions (3). Under pathological conditions, retinal microglia participate in potentiating neurodegeneration in diseases such as glaucoma, retinitis pigmentosa, age-related neurodegeneration, ischemic retinopathy, and diabetic retinopathy by producing proinflammatory neurotoxic cytokines or proangiogenic factors and by removing living neurons via phagocytosis (3).

Modeling the role of human microglia in the central nervous system has been a challenge because their differentiated state is environment dependent (4). When human microglia of the central nervous system (CNS) are extracted and cultured *in vitro* they de-differentiate and their transcriptional profiles are critically altered affecting expression of disease-linked genes. Therefore, the study of microglia *in situ* is key to understanding their role in development and disease. Mouse models provide a useful *in situ* environment, however, mouse microglia are distinct from human microglia as they lack the expression of many pathologically-linked genes present in humans (4). To understand the physiological and pathological roles of human microglia, it is important to study human microglia within a human retinal environment. There is currently no model of human retinal microglia in a mature human retinal niche that mimics form and function *in vivo*.

Recent work in cerebral organoids has found that resident microglia are present using standard culturing techniques that do not involve the ectopic addition of microglia or macrophage precursor cells (MPCs) (5). Using standard differentiation techniques for retinal organoids (ROs), microglia have not been found to be present in mature RO tissue (6, 7). Retinal organoids derived from human iPS cells (hiPSC) are differentiated through an ectodermal lineage and would not contain yolk sac-derived microglia. Recent protocols have been developed to generate microglia locally in RO cultures, however, these microglia have not been shown to integrate and function in mature retinal cell layers (8, 9). Other protocols have been developed by co-culturing early developmental ROs with separately differentiated iPSC-derived MPCs (10). These MPCs take up residence in developing ROs, however, microglial niches in mature ROs were not established.

During the course of development and in disease, microglia undergo morphological changes that correlate with their functional states (11). Ramified microglia which have fine, long processes are often referred to as “resting”, while ameboid microglia which have short processes with large soma are often referred to as “active” (11). Ramified microglia monitor neurons and are engaged in metabolite removal and clearance to keep homeostasis in the CNS, while ameboid microglia are engaged in synapse, axon, and cell engulfment with lysosomes and phagosomes (12, 13). At birth, retinal microglia are ameboid to remodel cells and synapses but become ramified as the retina matures (14, 15). Retinal injury, pathogen invasion, or neurodegeneration also lead to the formation of reactive ameboid microglia (11). During early development, yolk sac-derived microglia enter the retina via the circulatory system then infiltrate from the vitreal retina surface or the ciliary region in the periphery prior to vascularization (15, 16). A second microglia infiltration occurs from the optic disc or via blood vessels after vascularization. Microglia entry into the retina coincides with retinal neuron differentiation where they predominantly localize in the retinal synapse layers. In mice, almost all microglia initially localize to the nascent inner plexiform layer (IPL) when synapses begin to emerge in late embryonic stages (16, 17). Later, at postnatal day 9, microglia begin to localize in the newly formed outer plexiform layer (OPL). In a healthy retina, microglia and their processes remain predominantly localized to the inner retina and OPL and are largely absent in the outer nuclear layer (ONL) throughout adulthood (18).

To develop a 3D in vitro model of the retinal microglia niche in human retinal tissue we co-cultured mature ROs and MPCs, optimizing the parameters for successful integration of MPCs into ROs. We demonstrated that MPCs infiltrate and migrate to their proper anatomical locations, then develop a resting mature morphology and mature gene expression profile, corresponding to that observed in microglia *in vivo*. This model of mature induced human retinal microglia (iMGs) can be applied to future studies of microglial function in retinal disease.

## Results

ROs and MPCs were differentiated from a common clonal line of hiPSCs derived from a single donor. In our first attempts to integrate MPCs and ROs, we used ROs that were cultured for a minimum of 26 weeks, when retinal cells have matured well past proliferation. To establish a successful co-culture protocol combining MPCs and ROs, we needed to determine compatible media conditions, the duration of co-culture, and the number of cells to maximize the survival of both cultures together. We first determined the appropriate balance of culture media to allow for the health of both cell types in a single culture dish. MPCs are derived from differentiated embryoid bodies (EBs), called “Factories”, cultured in Factory media, whereas mature ROs are maintained in mature RO maintenance media. Both medias are incompatible with the differentiation and survival of the other cell type. However, a gradual shift from predominantly Factory media to pure RO media allowed for the survival of both cell types (Fig 1a). We added MPCs to ROs in an initial co-culture media of 3:1 Factory media:RO media. In subsequent days the media mixture was changed to 1:1, 1:3, then pure RO media (**Fig 1a**). During the first week of co-culture, we also maintained a concentration of M-CSF at 100 ng/mL to ensure the survival of MPCs until their integration into the ROs. During that time MPCs visibly adhered to the surface of the ROs. We found that media tapering and maintenance of 100 ng/ml M-CSF for one week were critical to MPC survival and integration. MPCs that were added to ROs in pure RO media or in the absence of M-CSF failed to adhere to the RO surface, and ROs that were maintained in a Factory/RO media mixture began to visibly disintegrate and die by 2 weeks. Residual free-floating MPCs were allowed in the co-culture for up to a week, then ROs with attached MPCs were completely removed and cultured in a clean well (**Fig 1a**). We found that the maximum saturation of MPCs in RO cultures was 1×10^6^ MPCs into a 6 well culture dish of 20-30 mature ROs. The addition of 2×10^6^ MPCs or greater resulted in death and degradation of the ROs as seen by disintegration of the spheroid. Successful co-culture conditions led to the the presence of CD45 and IBA1 positive cells within the RO retinal layers and in the RO lumen in as little as 2 weeks (**Fig 1b**). Next, we characterized the time course of the MPCs populating the ROs and found that the MPC population in ROs increased up to 4 weeks following the initial co-culture, post co-culture (PC), and remained stable up to 8 weeks PC, where the mean number of MPCs was 6.0±2.1/mm (**Fig 1c**). CD45 and CD11B positive cells accounted 0.5±0.1% of total RO cells (**Fig 1d, e**). This is comparable to 0.2-0.93% reported for adult mouse or rat retina (16, 19).

**Figure 1.**
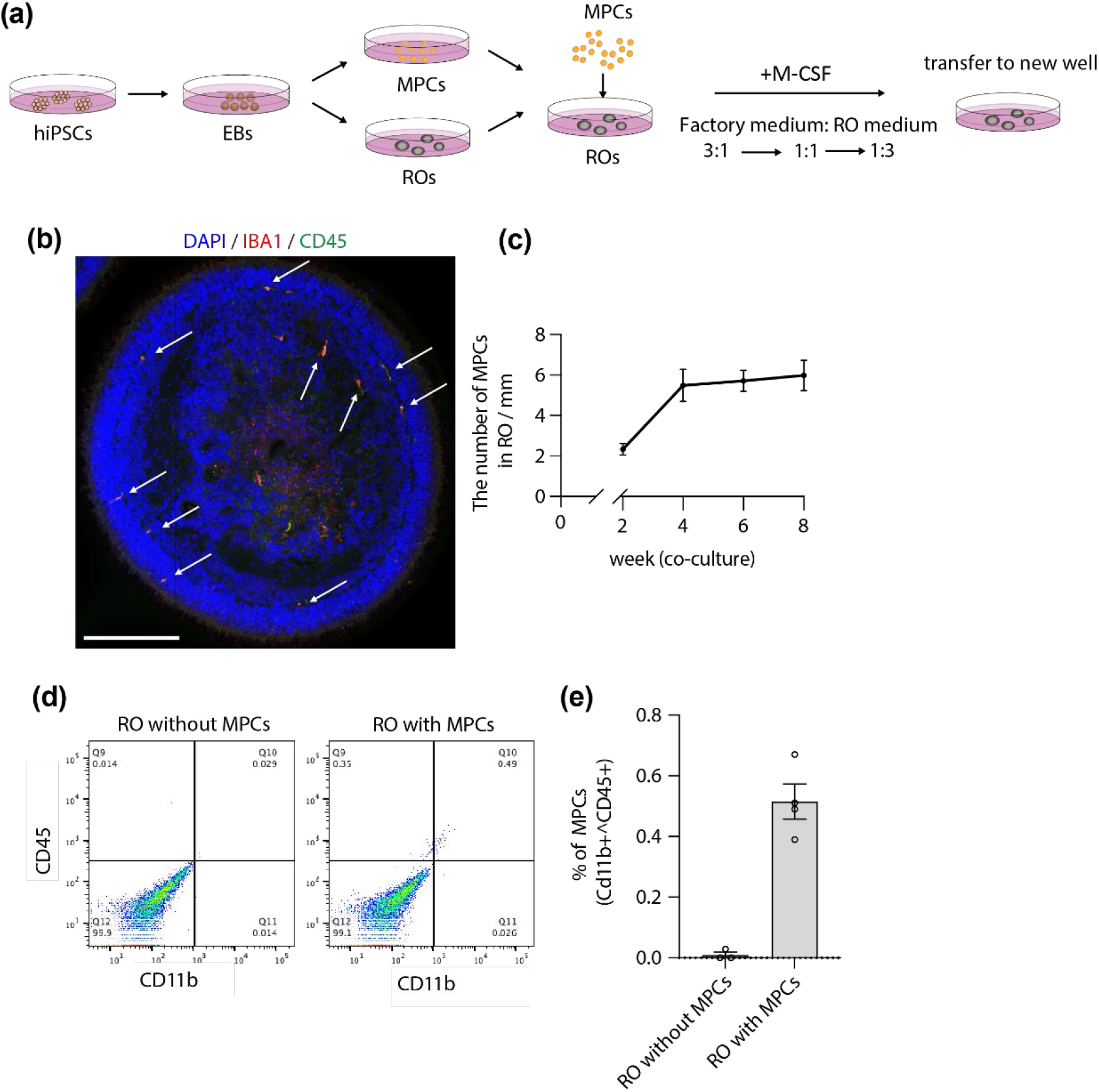
Optimized co-culture of MPCs and ROs allows for integration of CD11b/IBA1/CD45-positive MPCs into mature ROs. (a) Schematic illustrating the differentiation of MPCs and ROs from hiPSCs and subsequent co-culture parameters including media tapering and transfer of ROs with integrated MPCs to a clean culture well one week after MPC addition. (b) Representative confocal image of CD45+ (green) and IBA1+ (red) MPC within a mature RO localized to retinal layers and within the lumen. Scale bar = 200 µm. (c) Time course of MPC integration into ROs showed MPC population increased until 4 weeks then remained stable until week 8 (n=8-10). Data are shown as mean ± SEM. (d) Representative flow sorting chart showing a population of CD11b+ and CD45+ MPC only present in RO co-cultured with MPC (6-week PC). (e) CD11b+ and CD45+ MPC comprise 0.5±0.1% of total cells in ROs co-cultured with MPC as measured by flow sorting (n=4). Data are mean ± SEM.

We next wanted to determine if the age of ROs and MPCs would alter the efficiency of integration. Since our aim was to generate an integrated model of iMG in mature RO we restricted the minimum age of ROs to 20 weeks. At this age, ROs have completed the vast majority of their proliferative stage and most cells are post-mitotic and undergoing maturation (6). Co-culturing following proliferation prevents interference with altering RO differentiation, as the Factory media contains multiple potent growth factors. When we tested the efficiency of MPC integration into ROs aged 20-35 weeks, we observed that the age of post-mitotic ROs had no effect on the number of MPCs that integrated (**Supp Fig 1a**). Similarly, we tested efficiency of MPCs derived from MPC-producing EB “Factories” between 6 and 12 weeks old and found that Factory age, within that range, had no effect on MPC integration (**Supp Fig 1b**). This demonstrates that the integration of MPCs into ROs is a robust event that can occur under a wide range of cellular parameters. We further demonstrated that we could successfully replicate MPC-RO integration in a separate iPSC line from a separate donor showing that the protocol is repeatable (**Fig 2a**). We next tested to see if iMG-RO integration is compatible between MPCs and ROs differentiated from separate donor iPSCs. When we co-cultured from MPCs donor #2 (male) with ROs from donor #1 (female) we found that MPCs from donor #2 successfully integrated into donor #1 ROs (**Supp Fig 1c**).

**Figure 2.**
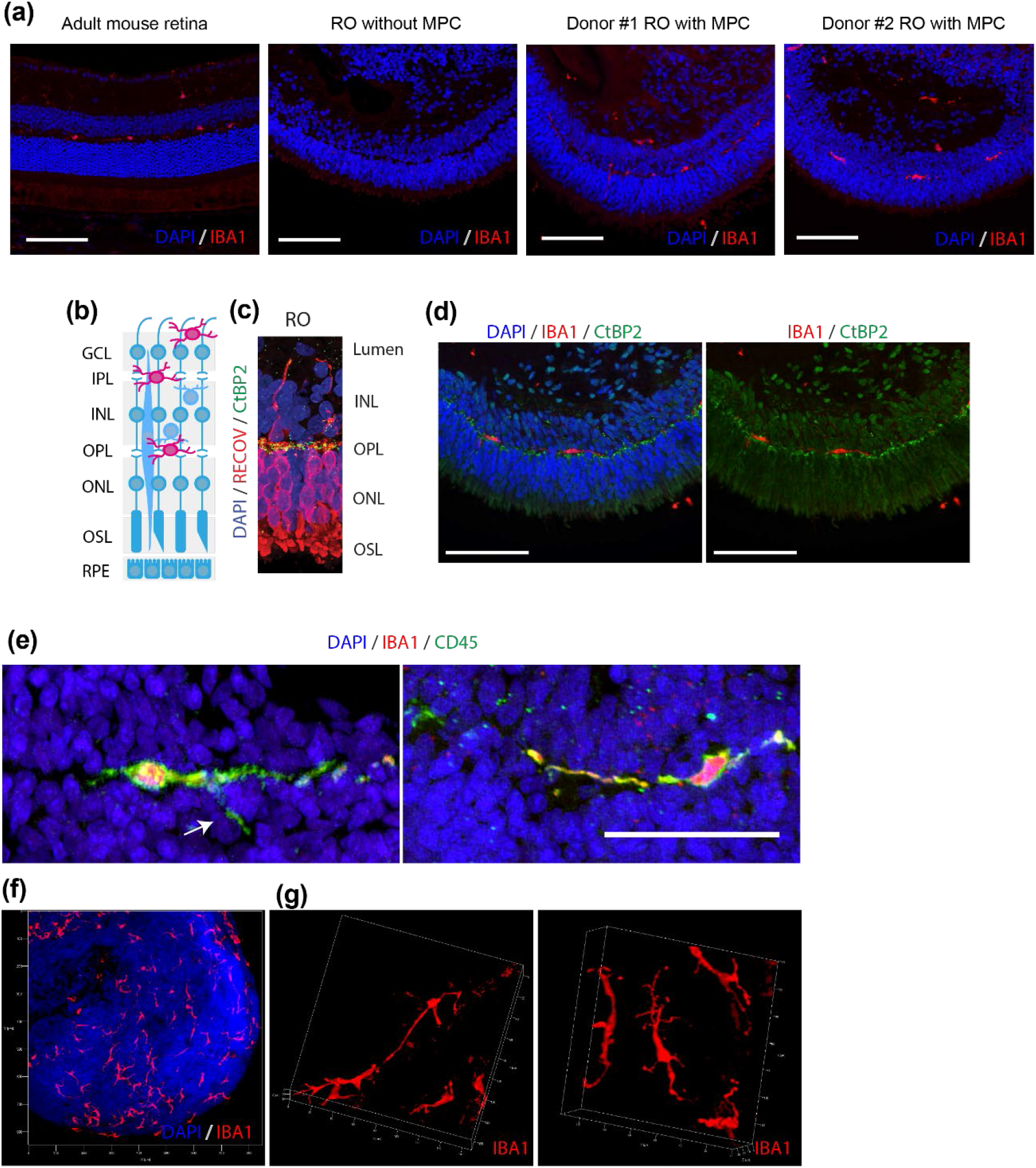
MPC localize to the OPL and the lumen within RO. (a) Panel 1(left): confocal image showing IBA1+ (red) microglia localized to the OPL and IPL in an adult mouse retina (10-week old C57/B6J). Panel 2 (middle left): 28-week RO without MPC co-culture showing absence of IBA1+ MPC. Panel 3 and 4 (Middle right, right): 31-week ROs from two different donors co-cultured with MPC for 6 weeks showing presence of IBA1+ MPC within synaptic layers of RO. Scale bars = 100 µm. (b) Cartoon showing the order and location of the various layers of the human retina. (c) Confocal image of RO showing stratification of retinal layers similar to human retina: photoreceptors (OSL/ONL: Recoverin, red), synaptic layer (OPL: CtBP2, green), second order neurons (INL: DAPI, blue), and a lumen. RPE: retinal pigment epithelium, OSL: outer segment layer, ONL: outer nuclear layer, OPL: outer plexiform layer, INL: inner nuclear layer, IPL: inner plexiform layer, GCL: ganglion cell layer. (d) MPC (IBA1, red) within mature RO localize to the OPL (CtBP2, green). Scale bars = 100 µm. (e) MPC (CD45, green) in the OPL were observed in the resting state with fine, long processes. Scale bars = 50 µm. White arrow indicates MPC process extending into the ONL. (f and g) 3D images of IBA1+ MPCs in whole mount ROs at 10x (e) and 40x (f) showing the architecture of the MPC processes in ROs.

Once we established a protocol for successful MPC-RO integration, we next characterized where in the RO the MPCs resided and if they matured to resting microglia. In healthy human and mouse retinas, microglia localize to the outer and inner plexiform layers (OPL and IPL respectively) (**Fig 2a,b**) where they assume a resting state characterized by ramified and elongated processes and suppressed expression of inflammatory cytokines (18, 20). Mature RO cultures have stratified retinal layers similar to the human retina, including an outer nuclear layer (ONL) comprised solely of the nuclei of photoreceptors, an OPL comprised of synaptic junctions, and an inner nuclear layer (INL) comprised of the nuclei of second order retinal neurons and Müller glia (21) (**Fig 2b,c**). The OPL is very well conserved in the RO (22), however, the IPL and the ganglion cell layer are less well-represented in ROs because retinal ganglion cells fail to survive in great numbers to mature ages, therefore, ROs lack a pronounced layer of synaptic junctions inside of the INL (23) (**Fig 2c**). We hence refer to the layer inside of the RO INL as the lumen. After 6 weeks co-culture, the MPCs are conspicuously absent from the ONL and INL which is very similar to normal *in vivo* adult mouse retina (**Fig 2a**). We found that IBA1 positive MPCs in ROs are localized mainly in the OPL, as defined by the ribbon synapse marker CtBP2, and the lumen (**Fig 2c,d**). Morphologically, MPCs in the RO OPL had fine long processes along the OPL band, which is indicative of microglia in a resting state (**Fig 2a-e**). 3D images of the MPCs in ROs show that their processes have complex branching and do not overlap with neighboring MPCs, similar to their in vivo counterparts (**Fig 2f,g**). However, some MPCs had processes branching into the ONL, which is not frequently observed *in vivo* (**Fig 2e, Fig3a**, white arrows).

We next characterized the timeline in which MPCs migrate to the different layers of ROs. At 2 weeks PC, we observed that the majority of MPCs initially reside in the INL and on the lumen side of the INL (**Fig 3a, b**). At the early time points MPCs have a spherical ameboid morphology indicative of an activated state which is conducive to migrating microglia (**Fig 3a**). The MPC population increases until week 4, where numbers are maintained (**Fig 3a, b**). Initially, at 2 weeks, few MPCs are observed in the OPL, however we observe the progressive accumulation of MPCs to the OPL that continues up to 6 weeks and stabilizes by week 8 (**Fig 3a, b**). The population of the OPL at later weeks correlates with MPCs developing elongated processes indicative of a mature microglial resting state (**Fig 3a**).

**Figure 3.**
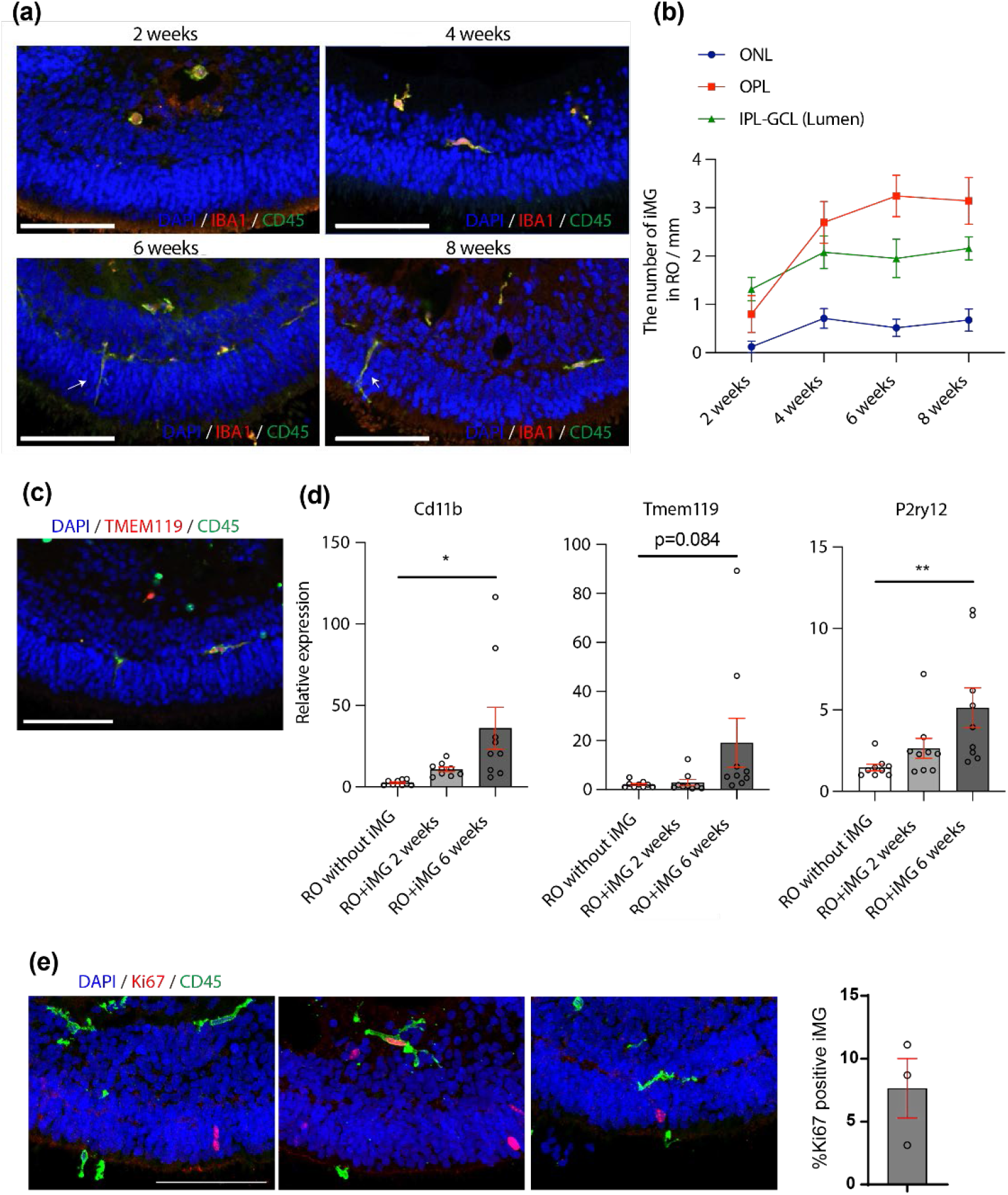
MPC in RO achieve a mature resting state by 6 weeks PC. (a) Representative confocal images showing MPC (CD45+, green and IBA1+, red) within lumen (GCL+IPL) of RO display spherical activated morphology at 2 weeks PC. By 6 weeks PC, the majority of MPC migrate to the OPL and mature to a resting state with long processes. ONL: outer nuclear layer, OPL: outer plexiform layer, IPL: inner plexiform layer, GCL: ganglion cell layer. White arrow indicates MPC process extending into the ONL. (b) Time course showing the migration of MPC to synaptic layers of RO from 2 weeks to 8 weeks PC. Scale bar = 100 µm (c) Representative confocal image showing at 6 weeks PC, iMGs (CD45+, green) in the OPL express TMEM119 (red) Scale bar = 100 µm. (d) Expression of pan-leukocyte marker CD11b and mature resting microglia markers TMEM119 and P2ry12 increase from week 2 to week 6 PC, determined by qPCR (n=9 each). Data are shown as mean ± SEM. \**p*<0.05, \*\**p*<0.01. (e) Representative confocal images showing iMG (CD45, green) in RO and dividing cells (Ki67, red) Scale bar = 100 µm. Panel 2 shows a dividing iMG. Quantification of Ki67+ iMG showed 5.1±3.3% of iMG in RO were dividing (n=3). Data are shown as mean ± SEM.

To determine whether MPCs have assumed a mature developmental state, we used the mature resting microglia marker TMEM119. We observed by immunohistochemistry that MPCs become positive for TMEM119 in the OPL at 6 weeks PC (**Fig 3c**). We confirmed through qPCR that the expression of TMEM119 and additional microglia markers pan-leukocyte marker, CD11b, and mature resting microglia marker, P2ry12, increase by week 6 PC coinciding with MPCs assuming a mature inactive morphology (**Fig 3a,d**). The collective observation of the integration of MPCs into the appropriate anatomical retinal layers with the development of a ramified microglial morphology and the presence of mature microglial expression profile validates that MPCs have assumed a mature microglial identity in a resting state within ROs by week 6 PC. We hence refer to MPCs in ROs as iMGs.

Next, we determined if iMGs populate the ROs through migration alone or if the progressive increase in the number of iMGs was due to proliferation within the RO following an initial migration. We used the proliferation marker Ki67 to determine that 5.1±3.3 % of iMG present in RO tissue are in a proliferative state. Under physiological condition, about 10% of microglia are dividing in the mouse retina, which is consistent with the percentage of iMG we observe dividing in ROs (24, 25).

The progressive morphological change of iMG from a spherical to an elongated shape in ROs suggests that iMGs transition from an active state at week 2 to an inactive state by week 6 once they take up a stable residence in the OPL. To validate this transition, we characterized the expression levels of MG active state markers in iMG-ROs at 2- and 6-weeks PC. We found that pro-inflammatory cytokines IL1b, TNFa, and IL6 were highly elevated at 2 weeks PC to 600x, 50x, and 18x that of control respectively, then return to control levels by 6 weeks PC (**Fig 4a-c**). The activation at week 2 PC and suppression by week 6 of pro-inflammatory cytokines also coincides with the increase of anti-inflammatory cytokines IL10 and TGFb1, which are significantly increased to 60x and 1.5x control, respectively, by 6 weeks PC (**Fig 4d,f**). We also noticed a slight but insignificant increase in BDNF expression by week 6 PC, indicating potentially beneficial physiological support of iMG following migration and inactivation (**Fig 4e**). Finally, we performed TUNEL staining to observe cell death proximal to integrated iMGs at 6 weeks PC (**Supp Fig 2**). We did not observe dying cells directly adjacent to iMGs integrated into the RO layers, confirming that iMGs were in fact inactive and not attacking the host cells. Collectively our data has shown that following 6 weeks, integrated MPCs achieve a stable resting microglial state predominantly in the OPL of the RO layers, establishing a mature outer retinal-microglial niche.

**Figure 4.**
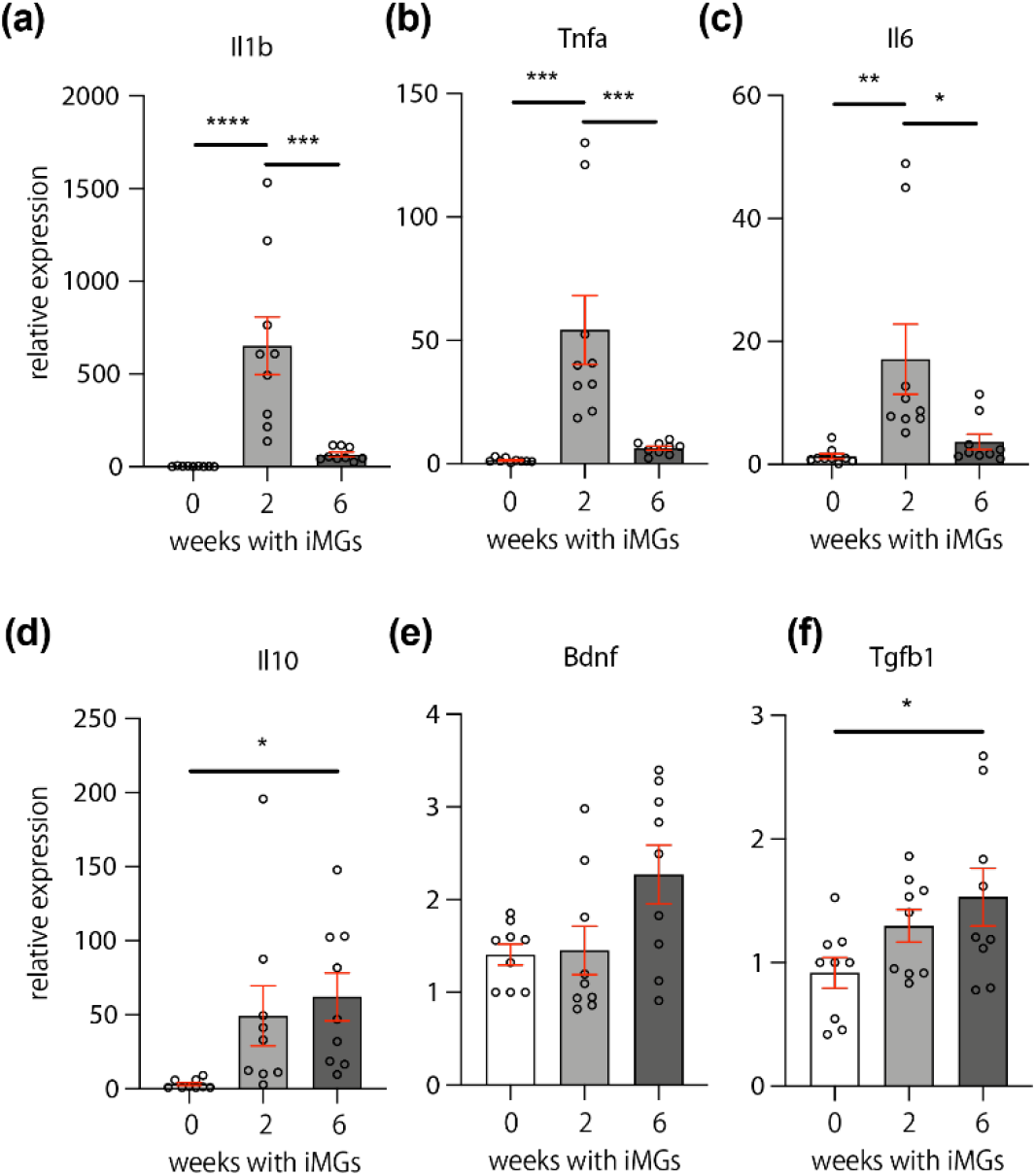
iMG in RO express pro-inflammatory cytokines at 2 weeks PC and anti-inflammatory cytokines at 6 weeks PC. Relative expression of pro-inflammatory cytokines (a) Il1b, (b) TNFa, and (c) Il6 were significantly elevated in RO with iMG at 2 weeks PC compared to control RO and RO with iMG at 6 weeks PC as measured by qPCR. At week 6 PC, relative expression of anti-inflammatory cytokines (d) Il10, (e) Bdnf, and (f) Tgfb1 are elevated compared to control RO and RO with iMG at 2 weeks PC as measured by qPCR. Data are shown as mean ± SEM. \**p*<0.05, \*\**p*<0.1, \*\*\**p*<0.01, \*\*\**p*<0.001.

## Discussion

Microglia play a crucial role in the development, physiology, and function of the human retina, however current models of mature retinal organoids do not contain a microglial niche. Here we show, for the first time, the successful integration of MPCs into mature ROs, thereby generating a more complete retinal tissue model containing mature iMG. This model provides an important technical advance for the study of retinal microglia directly in human tissue.

The development of *in vitro* models to study human retinal microglia has been a challenge for the field. Human microglia, whether iPSC-derived or isolated from fresh tissue, that are grown in simple monocultures lack the environmental complexity required for them to mature to a state that resembles their identity and functions *in vivo*. Animal models have provided powerful surrogates; however, they have fundamental differences that hinder their ability in microglia to faithfully model human disease and development (4). In our co-culture model, we optimize and characterize the conversion of MPCs to resident mature microglia in ROs. We show that MPCs in co-culture initially migrate into the RO where they cycle through an activated phase, characterized by round morphology and elevated expression of inflammatory cytokines, at around 2 weeks PC. By 6 weeks PC, they establish a niche in the OPL where they mature into resting microglia, adopting morphology characterized by small cell bodies and long branching processes that stretch along the synaptic rich plain. We further validate that in the OPL, microglia express cell specific gene markers characteristic of mature microglia. Using this technique, we recreate a mature retinal microglial environmental niche that allows for the modeling of retinal microglial-specific functions directly in human tissue. Further, it would be possible to develop this model into a human retinal disease model using diseased iPSCs or culturing pathologic conditions.

Microglia are important immune cells of the retina that play a central role in mediating pathological degeneration in diseases such as glaucoma, retinitis pigmentosa, age-related neurodegeneration, ischemic retinopathy and diabetic retinopathy (3). This co-culture system will be useful for understanding the pathogenesis of retinal diseases involving pathological activity of retinal microglia and provide a platform for future drug discovery. As a promising sign of reproducibility, we were able to replicate this co-culture protocol in two separate iPSC lines from two individual donors. We also found that co-cultured iMGs and ROs from separate donors were compatible. This versatility will allow for the integration of iMG and RO with different genetic backgrounds to isolate cell-specific genetic effects. This feature will greatly improve the ability to study human-specific pathways and pharmacological responses. A human retinal environment is necessary to understand features unique to human microglia and their pathological reaction. For example, HTRA1, C2, and C3 which are genetic risk variant for age-related macular degeneration are expressed in human microglia, but not in mice microglia (4).

Although the co-culture model presented here is the first to develop a mature retinal microglial niche, this model has several limitations. One major limitation is that mature organoids are predominantly models of the outer retina and lack well organized and stable inner retinal features. Retinal ganglion cells do not survive in large numbers at later stages of retinal organoid maturity making the inner plexiform layer (IPL) nearly nonexistent. In the retina, the IPL and retinal ganglion cell layer are critical microglial niches. Future efforts to improve the survival of the inner retinal layers will be critical for further integration of microglia. Another important limitation is the relatively low integration of microglial populations. Despite changing multiple variables, we were unable to achieve saturation of the OPL with microglia. Due to the low rate of dividing microglia, it is unlikely that extended culture times would allow for larger populations. RO models also lack functional vasculature which are critical to a complete microglial niche. Future developments that also integrate a vascular network may also facilitate the integration of microglia into ROs.

In our efforts to define the culturing parameters of the protocol we observed that variables such as the age of the mature organoid and the age of the MPC Factory culture did not have noticeable effects on the number of MPCs that integrated into ROs, suggesting that ROs and MPCs have a long period of receptivity to integration. Of the variables under our control, we found that the quality of the MPC Factory culture was the biggest indicator of successful integration. Fresh trophic factors and media preparations were required for optimal MPC Factory conditions. We also observed that the initial mixing of Factory and RO culture media with a gradual change to RO media and M-CSF supplementation was essential for MPCs to remain healthy when added to ROs. MPCs from the Factory culture are highly unstable and sensitive to changes in media composition. Great care should be taken in the first co-culture step to allow for them to remain MPCs and integrate into ROs for further maturation.

Organoid cultures provide great promise for the study of complex human tissues. Until recently, they have been restricted to cell types derived from a limited number of progenitors from single germ layers, lacking the true complexity of organs. By integrating yolk sac-derived microglia into neural ectoderm ROs, we have made an important technical advance in the complexity of retinal organoid models.

## Materials and Methods

### Cell and cell culture

#### Stem cells

hiPSC lines were derived from peripheral blood mononuclear cells from a female (donor #1) and male (donor #2). Reprogramming was performed using sendai virus for reprogramming factor delivery. Donor #1 hiPSCs were derived by the Harvard iPS core facility and donor #2 hiPSCs were derived by the Salk iPSC core facility. All cell lines were obtained with verified normal karyotype and contamination-free. hiPSC were maintained on Cultrex (R&D Systems, 3432-005-01) coated plates with mTeSR+ medium (STEMCELL Technologies, 100-0276). Cells were passaged every 3-4 days at approximately 80% confluence. Colonies containing clearly visible differentiated cells were marked and mechanically removed before passaging.

#### Microglia precursor cell differentiation

MPCs were generated as previously described (26, 27). The embryoid bodies (EBs) are formed using Aggrewells (STEMCELL Technologies, 34811), cultured with bone morphogenetic protein 4 (BMP4, Peprotech, 120-05), vascular endothelial growth factor (VEGF, Peprotech, 100-20), and stem cell factor (SCF, Miltenyi, 130-093-991), then plated into T175 flasks with Interleukin-3 (IL3, R&D Systems, 203-IL-050) and macrophage colony-stimulating factor (M-CSF, ThermoFisher, 203-IL-050). After 4 weeks, MPCs emerged into the supernatant. MPCs were harvested for co-culture between 6-12 weeks. It was previously revealed that their ontogeny is MYB-independent primitive myeloid cells, which is the same ontogeny as microglia (28).

#### Retinal organoid differentiation

To make retinal organoids, embryoid bodies were formed (day 0) using Aggrewells (STEMCELL Technologies, 34811) in Neural Induction Medium and then plated (day 7) onto Matrigel growth factor reduced (Corning, 47743-718) coated plates. Media was switched to Retinal Differentiation Medium (RDM) on day 16. On day 28, plated EBs were fragmented using a sterile scalpel (McKesson, 801453), scraped with a P1000 pipette, collected, and transferred to suspension culture on an orbital shaker. At week 8, mature organoids were transferred to a bioreactor, and the medium was changed to Retinal Differentiation Media with 10% FBS, 100 µM taurine, and 2 mM Glutamax (RDM+). The remainder of the differentiation protocol is as previously published (6, 29). Organoids were transferred from the bioreactor at week 17 and plated in low attachment 6 well plates.

#### RO-MPC co-culture

Harvesting of MPCs was as previously described (22, 23). Supernatant from the Factory containing MPCs was passed through a cell strainer (Corning, 431750). Cells were spun at 400G, resuspended in fresh Factory Media, counted, and 10^6^ MPCs were added to 20-30 mature retinal organoids in a low attachment 6 well dish in 3:1 Factory Media to RDM with M-CSF at 100 ng/mL (day 0). In subsequent days, the media mixture was changed to 1:1 with M-CSF, then 1:3 with M-CSF, taking care not to aspirate any iMG that hadn’t yet adhered to the exterior of the ROs. On day 4, the media was changed to RDM+ with M-CSF until day 7 when the iMG-RO co-culture was transferred to a clean well. Upon transfer, the co-culture was maintained in RDM+ without M-CSF, resulting in the loss and detachment of MPCs from the surface of ROs.

### Immunohistochemistry

Retinal organoids were placed in 4% paraformaldehyde (PFA) in PBS for 15 minutes. Organoids were washed in PBS then submerged in 20% sucrose in PBS overnight. Tissues were embedded in Tissue-Tek OCT compound (Sakura Finetek; Torrance, CA) and frozen at −80°C. Cryosectioning was done at 14 µm and sections were mounted on glass polylysine coated slides. Prior to primary antibody addition, sections were blocked with 2% bovine serum albumin and 0.1% Trtion X-100 in PBS for 30 minutes. Primary antibodies were added to samples at 4°C and incubated overnight. Primary antibodies targeting IBA1 (1:500; FUJIFILM, 019-19741), CtBP2 (1:200, BD Biosciences, 612044), Ki67 (1:100, Invitrogen, PA5-16446), Recoverin (1:500, Millipore, AB5585), TMEM119 (1:500, Sigma, HPA051870) and CD45 (1:200, BD Biosciences, 555480) were used. Sections were incubated with species-specific secondary antibodies (1:1000) for 1 hour at RT then washed in PBS and mounted in mounting media (VectaShield, H-1000). All images were acquired with a confocal laser scanning microscope (LSM 710, Zeiss) and processed with the ZEN 2010 software (Zeiss).

#### Mouse retina IHC

Dissected eyes were incubated in 4% PFA for 1 hour. Eyes were washed in PBS then submerged in 20% sucrose in PBS overnigh and embedded in Tissue-Tek OCT compound for cryo-sectioning and frozen at −80°C. 14-μm retinal sections were washed in PBS and incubated in blocking buffer (PBS with 2% bovine serum albumin, and 0.1% Triton X-100) for 2 hours at 4°C, following by an overnight incubation with primary antibodies targeting IBA1 (1:500; FUJIFILM, 019-19741) in blocking buffer at 4°C. Sections were incubated with species-specific secondary antibodies (1:1000) for 1 hour at RT then washed in PBS and mounted in mounting media (VectaShield, H-1000). All images were acquired with a confocal laser scanning microscope (LSM 710, Zeiss) and processed with the ZEN 2010 software (Zeiss).

### 2.4 Flowcytometry

A postnatal neural dissociation kit (Miltenyi, 130–092-628) was used to prepare a single cell suspension from retinal organoids co-cultured with MPCs. Cells were centrifuged at 150 g for 5 min at 4°C. The digested tissue was resuspended in 100 μL of 4% FBS in PBS containing an FITC antibody to CD11b (1:100; BioLegend, 101206) and PE antibody to CD45(1:100; BD Biosciences) and incubated for 20 min on ice. The cells were washed and suspended with 1 mL of 4% FBS/PBS containing DAPI (1:2000; Thermo Fisher Scientific, 62248) and DRAQ5 (1:5000; Cell signaling, 40845) for exclusion of dead cells and debris.

### 2.5 RNA isolation and real-time PCR

Retinal organoids co-cultured with MPCs were collected in 500 μL of Trizol and total RNA was isolated using a RNeasy micro kit (QIAGEN) according to manufacturer’s instructions, and reverse transcribed using Maxima First Strand cDNA Synthesis Kit for RT-qPCR (Thermo Scientific). qPCR was performed using Power-up SYBR™ Green PCR Master Mix (Thermo Fisher Scientific) and primers on a Quantstudio 5 Real-Time PCR System (Thermo Fisher Scientific). β-actin (*Actb*) was used as the reference gene for all experiments. Levels of mRNA expression were normalized to those in controls as determined using the comparative CT (ΔΔCT) method. Primer sequences are listed in Table S1.

### Quantification and statistical analysis

All statistical tests were performed in GraphPad Prism v8 (GraphPad Software, Inc). Data comparisons between two groups were performed using unpaired two-tailed Student t-tests. Data comparisons between multiple groups were performed with one-way ANOVA with Tukey’s correction. Statistical tests used for each experiment are specified in the figure legends. Data are represented as mean ± SEM. A p value of p < 0.05 was considered significant.

### Study approval

All animal protocols were approved by the IACUC committee at The Scripps Research Institute, La Jolla, California, and all federal animal experimentation guidelines were adhered to.

## Acknowledgments

We thank the members of the Friedlander lab and our colleagues at the Lowy Medical Research Institute for many helpful discussions regarding the direction of this project and the data presented in this manuscript. We would like to thank TSRI’s Flow cytometry core for their excellent technical assistance, and TSRI’s animal vivarium staff for the excellent care of the animals used in this study. This work was supported by the Lowy Medical Research Institute and NIH grants EY11254 (to M.F.). A.U.-O. is supported by a fellowship from the Manpei Suzuki Diabetes Foundation and JSPS KAKENHI Grant Number 17K16984 and 21K09727.

We would like to thank the members of the Friedlander laboratories and our colleagues at the Lowy Medical Research Institute for many helpful discussions. This work was supported by the Lowy Medical Research Institute (M.F.) and NIH grants EY11254 (M.F.). A.U.-O. was supported by a fellowship from the Manpei Suzuki Diabetes Foundation and JSPS KAKENHI Grant Number 17K16984 and 21K09727.

## Data Availability Statement

The data that support the findings of this study are available from the corresponding author upon reasonable request.

## Figures

**Supplementary Figure 1.**
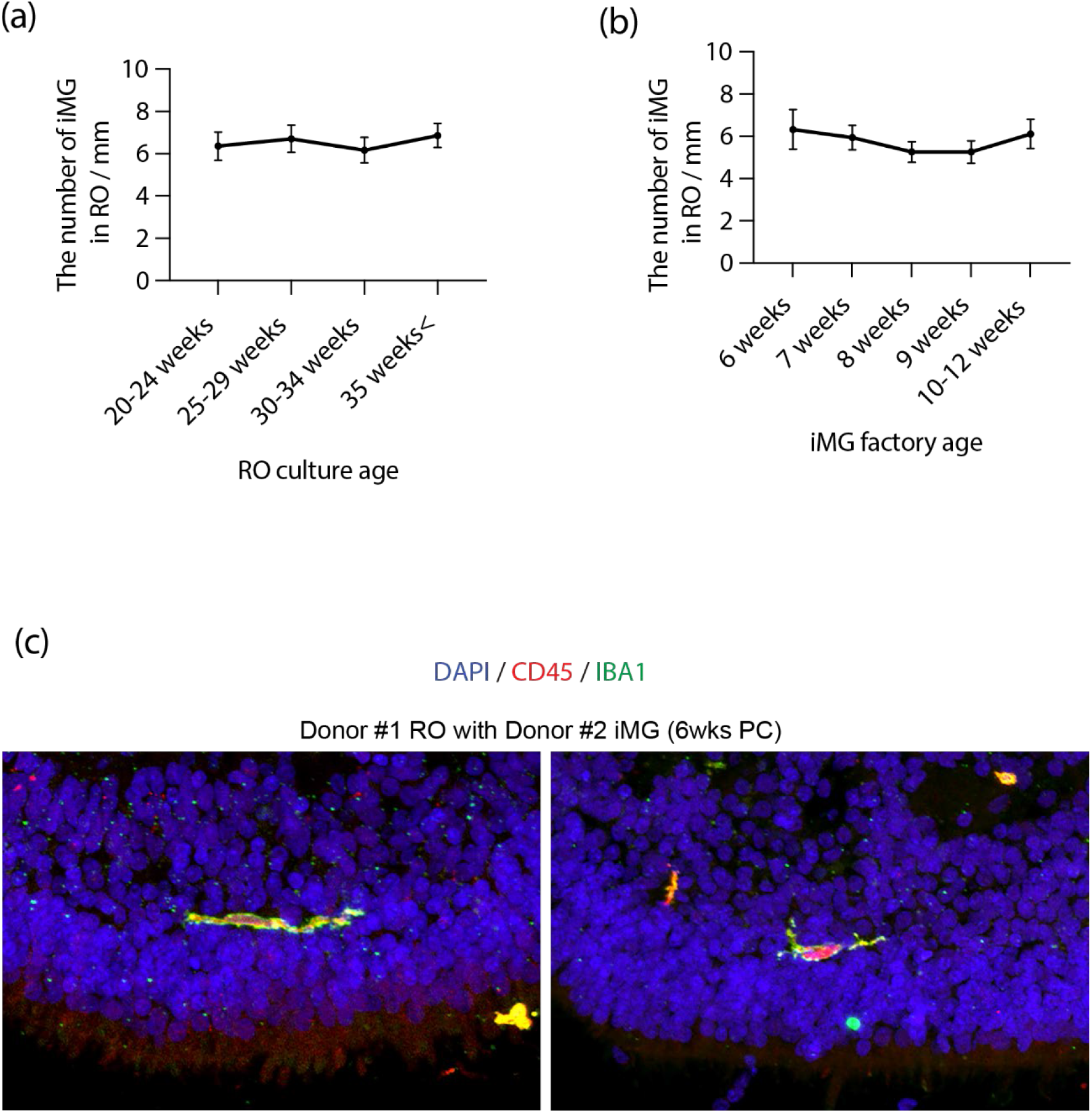
Efficiency of iMG and RO integration was unaffected by the age of mature RO, the age of the MPCs, or the cell line used for either cell type. (a) The number of MPCs present in mature ROs was unaffected by the age of the RO, between 20-35 weeks (n=8-10). Data are shown as mean ± SEM. (b) The number of MPCs present in mature ROs was unaffected by the age of the MPCs upon harvest from the Factory (n=8-10). Data are shown as mean ± SEM. (c) Representative confocal images showing that CD45+ and IBA1+ MPCs are localized to retinal layers when mixing MPCs from donor #2 with RO from donor #1.

**Supplementary Figure 2.**
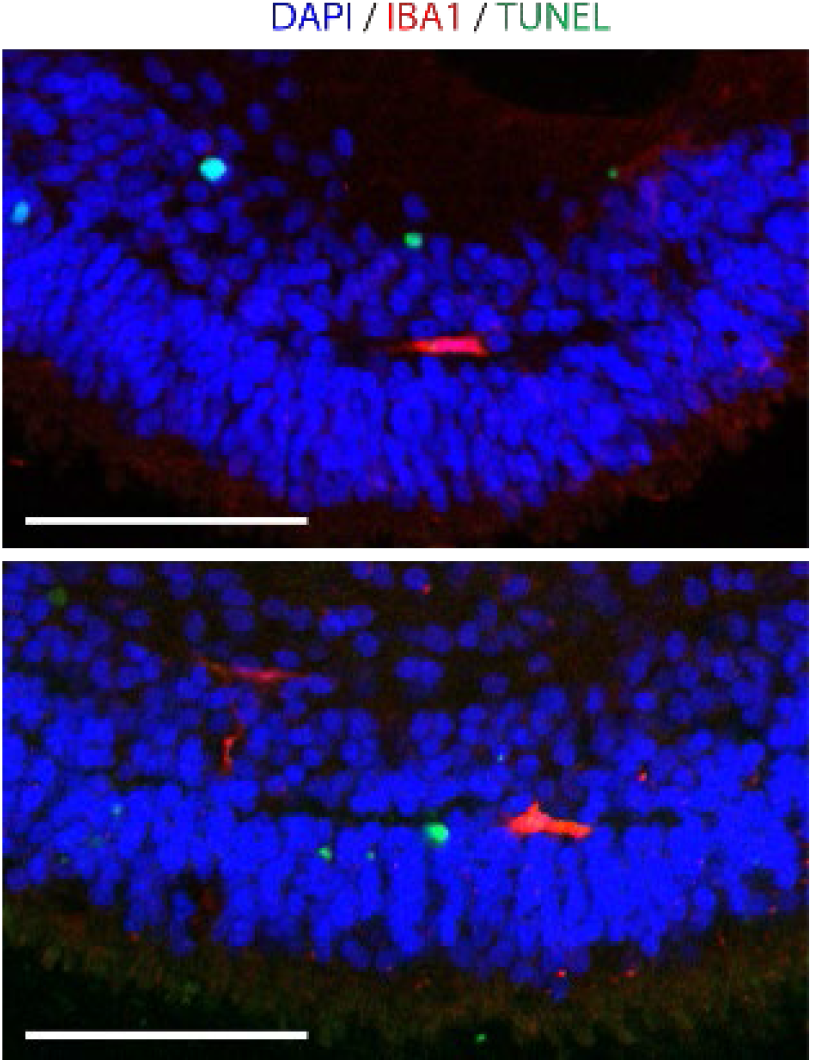
iMG integrated into RO layers are not attacking host cells. (a) Representative confocal images show iMG (CD45+, red) are not directly adjacent to dying cells (TUNEL, green).

